# Urban chemical stress disrupts cross-domain microbial networks in river sediments

**DOI:** 10.64898/2026.03.13.711587

**Authors:** Eduardo Acosta, Thomas Backhaus, Werner Brack, Pedro A. Inostroza

## Abstract

Freshwater sediments face increasing anthropogenic stress, yet their effects on cross-domain microbial interactions remain understudied. We analysed co-occurrence networks encompassing bacteria, protists, and fungi in sediments from two urban river systems in central Chile (Aconcagua and Maipo), using high throughput metabarcoding. Sites under multidimensional pollution stress exhibited fragmented networks, reduced cross-domain connectivity, and a predominance of positive correlations, consistent with stress-induced facilitation. Keystone taxa shifted, with Firmicutes, Ciliophora and Rozellomycota gaining prominence under stress. Conversely, reference sites displayed cohesive networks and balanced interaction types, driven by Proteobacteria, Ascomycota and Basidiomycota. Negative co-occurrences between protists and bacteria in contaminated zones suggest intensified competition or top-down trophic control. Our findings highlight the vulnerability of cross-domain microbial assemblages to urban pollution and identify specific metrics as candidate bioindicators for ecosystem integrity.

## 1. Introduction

The riverine microbiome in urban environments is shaped by multiple anthropogenic stressors, including exposure to organic micropollutants (OMPs) from treated and untreated wastewater, as well as agrochemicals runoff (Inostroza et al., 2025; Martínez-Santos et al., 2018; Verbücheln et al., 2025). The interactions of OMPs and the environmental microbiome in sediments have far-reaching consequences for ecosystem functioning (Burdon et al., 2020; Inostroza et al., 2025). Sediments serve as biogeochemical hotspots, driving nutrient cycling and pollutant degradation–through the activity of diverse, metabolically versatile microorganisms (Borton et al., 2025; Liu et al., 2023). Within these communities, microbial interactions underpin essential processes that influence water quality, ecosystem stability, and global biogeochemical fluxes (Battin et al., 2007; Borreca et al., 2024; Gao et al., 2022). Yet the increasing burden of environmental pollutants –particularly mixtures of OMPs–can disrupt microbial interactions and impair ecological functions, threatening ecosystem resilience (Gao et al., 2022; Inostroza et al., 2025). Understanding how these complex microbial assemblages respond to chemical pollution is thus central to predicting and managing the ecological risks of contamination in freshwater systems (Pelletier et al., 2020).

Deciphering microbial interactions in natural environments has long been a central goal in microbial ecology (Brock, 1966; Bungay, 1968). In this context, correlation-based co-occurrence networks are an emerging tool to infer potential ecological interactions from environmental DNA (eDNA) sequencing data (Srinivasan et al., 2024). In such networks, nodes represent microbial taxa, while edges indicate statistically significant co-occurrence or exclusion patterns, often interpreted as proxies for positive (e.g., facilitative) or negative (e.g., competitive) interactions (Faust and Raes, 2012). Key topological metrics–such as node degree, modularity, and betweenness centrality–reveal aspects of community organization and help identifying keystone taxa, which exert a disproportionate influence on ecosystem structure and function (Banerjee et al., 2018). For instance, taxa with high betweenness centrality, can serve as critical connectors, maintaining interaction pathways and buffering communities against structural collapse (Zamkovaya et al., 2021).

In polluted sediments, microbial networks are reorganized around specialist keystone taxa, particularly bacterial species adapted to metals and OMPs, including pharmaceuticals (Veloso et al., 2023). Furthermore, network analyses have recently been extended to explore cross-domain interactions, defined here as ecological associations among microorganisms from different phylogenetic domains, such as Bacteria, Archaea and Eukarya (e.g., protists and fungi) (Hou et al., 2024; Ng et al., 2025). These cross-domain associations are ecologically significant: protists regulate bacterial abundance through predation, while bacteria-fungi interactions shape organic matter decomposition through competitive or mutualistic pathways (Arndt et al., 2000; Romaní et al., 2006).

Emerging evidence suggests that environmental stressors–particularly OMPs–can rewire inter-phyla associations, reduce interaction strength, and weaken ecological resilience in bacterial communities (Gao et al., 2022). Notably, these patterns can be interpreted through the lens of the Stress Gradient Hypothesis (SGH), which postulates that increasing abiotic stress favours facilitative over competitive interactions (Bertness and Callaway, 1994). In microbial communities, this often manifest as a rise in positive co-occurrence correlations under stress, indicative of cooperation via syntrophy, resource sharing, or shared resistance mechanisms (Morris et al., 2013). For instance, networks in copper-polluted sediments show increased positive correlations and reduced modularity, suggesting stress-driven facilitation (Yuan et al., 2021). To date, only a handful of studies have employed the SGH in microbial network contexts–mainly focusing on prokaryotes or bacteria-fungi systems under abiotic stress such as drought or warming (Hammarlund and Harcombe, 2019; Hernandez et al., 2021). Although recent work involving soil protists, fungi, and bacteria demonstrates cross-domain network simplification under thermal stress (Zhou et al., 2021), there remains a notable gap in testing SGH across multi-domain networks in chemically polluted environments, particularly riverine sediments.

Applying the SGH to cross-domain microbial networks offers a novel lens through which to evaluate how OMP stress reshapes interaction patterns, ecological roles, and community resilience. Co-occurrence networks, in this context, can reveal domain-specific shifts in keystone taxa–nodes that integrate trophic or metabolic functions across microbiomes. The emergence or loss of these central nodes under pollution stress may signal ecological thresholds of collapse or adaptive reorganization, serving as potential early-warning indicators (Veloso et al., 2023). This perspective is especially relevant for sediment ecosystems, where microbial communities serve as biogeochemical keystones but remain largely underexplored in terms of species interactions.

To address this gap, we assessed microbial network structure and stability in the sediments of two river basins in central Chile, integrating pollution profiles–including nutrient and OMP levels in the overlaying water column. Specifically, we examined: i) the balance of positive and negative correlations in microbial networks across sites to infer shifts in interaction types (e.g., facilitation versus competition) among bacteria (prokaryotes), protists, and fungi (microeukaryotes) in response to pollution gradients (SGH); ii) network modularity as a proxy for ecological fragmentation and resilience; and iii) keystone taxa through betweenness centrality to identify cross-domain microbial integrators that maintain community connectivity under environmental stress. These basins offer a natural gradient of OMP contamination and have been used as model systems for studying pollution-driven ecological change in understudied river systems in South America (Cortés-Miranda et al., 2024; González-Rojas et al., 2025; Inostroza et al., 2024; Soriano et al., 2024).

## 2. Methods

### 2.1. Selection of study sites

The Aconcagua River (7,338 km^2^) and the Maipo River (4,843 km^2^) are two major river basins in central Chile. Both rivers play a crucial role in supporting the region’s socio-economic activities, particularly agriculture, which dominates land use within their basins(Soriano et al., 2024). In addition, both rivers receive effluents from multiple wastewater treatments plants (WWTPs), making them representative systems for studying the combined impacts of human activities on freshwater ecosystems (Inostroza et al., 2025, 2024; Soriano et al., 2024).

Seven sampling sites were selected across these river systems: Three in the Aconcagua River and four in the Maipo River (Supplementary Figure 1). Site selection was designed to represent a gradient of OMP impact–from relatively pristine conditions (reference sites) to sites influenced by urbanization and wastewater discharge (urban and rural sites). In the Aconcagua River, three sites included: ARS1, a reference site located in a pre-Andean upstream area with low anthropogenic influence, distant from major cities; AR2, situated within a densely populated area and subject to high urban pressure; AR3, a study site located in a rural area. In the Maipo River, four sites included: MRS1, a reference site located in a pre-Andean upstream area and isolated from major urban centres; MT1, located upstream a municipal WWTP within an urban setting; MT2, located downstream of the same WWTP, thus representing a site under direct influence of treated urban effluents; MR1, a rural study site. Detailed location of the samples sites is provided in Supplementary Table 1.

### 2.2. Environmental characterisation of Aconcagua and Maipo Rivers

At each study site, *in situ* measurements of physicochemical parameters in the water column, including pH, conductivity, temperature and dissolved oxygen were performed using a multiparameter probe (Hannah model HI 9829, United Kingdom) or HOBO data loggers (Onset, USA). To assess major nutrients, 15 mL surface water was filtered in triplicate through a 0.45 μm GF/F glass fibre filter and stored at 20 °C until laboratory analysis. Nitrite, nitrate and phosphate were measured using standard colorimetric methods with a segmented flow Seal Auto-analyser (Seal Analytical AA3).

Water samples for OMPs were collected at the same time as sediment samples were collected for eDNA metabarcoding. Details on OMP concentrations at each sampling site, along with comprehensive descriptions of sample collection, storage, extraction procedures, and LC-HRMS analysis, are described in Soriano et al., (2024). Thus, the OMP concentrations were retrieved from Soriano et al., (2024) and OMPs were grouped by chemical class, and their concentrations within each class were summed up to construct an environmental vector (Supplementary Table 2). Additionally, OMPs with antimicrobial properties (e.g., antibiotics) and those capable of disrupting prokaryotic and microeukaryotic photosynthesis were normalised to their effect concentrations applying the toxic unit approach (TU) described elsewhere (Inostroza et al., 2025; Verbücheln et al., 2025).

### 2.3. DNA-based characterization of microbial communities in riverine sediments

Surface sediment (top 0-5 cm) in direct contact with the water phase was collected using sterile 50 mL single-use tip-off syringes. For this, three biological replicates, were transferred into 5 mL sterile cryovials and stored in liquid nitrogen (Air Liquide Voyageur 12) until DNA extraction. The eDNA from sediments was extracted using the DNeasy PowerSoil Pro Kit (Qiagen, Germany), following the manufacturer’s instructions and suggested modifications (Inostroza et al., 2025). Bacterial 16S rRNA gene sequences were amplified using the V3-V4 primer pair (341F: CCTAYGGGRBGCASCAG; 806R: GGACTACNNGGGTATCTAAT) (Yu et al., 2005). Protist 18S rRNA gene sequences were amplified using the V9 primer pair (389F: GCCTCCCTCGCGCCATCAGXXXXXTTGTACACACCGCCC; 1510R: GCCTTGCCAGCCCGCTCAGCCTTCYGCAGGTTCACCTAC) (Amaral-Zettler et al., 2009). The fungal nuclear ribosomal internal transcribed spacer 2 (ITS2) was amplified using the primer pair (ITS3-2024F: GCATCGATGAAGAACGCAGC; ITS4-2409R: TCCTCCGCTTATTGATATGC) (Orgiazzi et al., 2012). The PCR amplification and amplicon sequencing were conducted by Novogene Co. (Cambridge, United Kingdom). All samples were sequenced using a 250-bp paired-end using a NovaSeq 6000 sequencer Sequencing System – Illumina (USA).

The raw forward and reverse sequences in “fastq” format obtained from the 16S rRNA gene (bacteria), the 18S rRNA gene (protists) and the ITS2 region of the ribosomal operon (fungi) were imported into the software QIIME2 v. 2024.5 (Bolyen et al., 2019). The primer sequences were removed from the libraries using the function “cutadapt trim-paired,” and sequences with an error rate higher than zero were discarded. After trimming, the sequences were dereplicated, denoised, and merged into amplicon sequence variants (ASVs), calling the function “denoise-paired” from the DADA2 plugin (Callahan et al., 2016). This step included chimera filtering of ASVs using the “isBimeraDenovo” method, which detects chimeras based on the consensus chimeric fraction across all samples. Singletons and doubleton were not removed from our datasets.

Taxonomic assignment was performed with the function “feature-classifier classify-consensus-vsearch” using a cut-off value of 0.8 and 97% identity with the SILVA database version 138.1 for bacteria. For the identification of protists, we used the Protist Ribosomal Reference (PR^2^) 4.14.0, amended with the V9 region of the 18 SSU rRNA gene sequences from the Heterotrophic Flagellate Collection Cologne (HFCC) (Schoenle et al., 2021). The classification of fungi was established through prior training of a classifier using the machine learning algorithm of the function “feature-classifier fit-classifier-naive-bayes” (Pedregosa et al., 2011) and the non-redundant version 10.0 of the UNITE+INSDC fungal ITS database as reference (Abarenkov et al., 2024). All sequences identified as bacteria, protists, or fungi were retained for downstream analysis. Sequences unclassified or assigned to non-target organisms were excluded from their respective datasets. To combine taxonomic groups for joint analyses, we merged feature tables and representative sequences using the the “feature-table merge” and “feature-table merge-seqs” functions. A phylogenetic tree was constructed using the “qiime phylogeny align-to-tree-mafft-fasttree” function. Additionally, we evaluated microbial community similarity across sampling sites using the Jaccard distance based on presence-absence data from the combined bacterial, protist and fungal ASVs. The resulting distance matrix was used to construct a hierarchical clustering dendrogram (cladogram), illustrating the sites grouping based on overall microbial community structure, including replicate samples per sites.

### 2.4. Statistics and reproducibility

From each study site, the following files were extracted using QIIME2: merged representative sequences, merged ASV frequency table, taxonomic assignments (for bacteria, protists and fungi), a phylogenetic tree of the merged sequences, and the associated metadata table. These files were imported into R version 4.3.0, using the “file2meco” package and processed using the “microeco” package (Liu et al., 2022, 2021). Co-occurrence networks were inferred in microeco using the “cal_networks() function with cor_method = “sparcc” (Friedman and Alm, 2012). ASVs with relative abundances below 0.001% were filtered out, and a null model with 100 bootstrap iterations was used to assess significance.

To evaluate the structure within each study site, modularity was calculated using the “cal_module” function with the “cluster_fast_greedy” method^41^. Networks attributes such as number of nodes, number of edges, and the proportion of positive and negative correlations were calculated using the “cal_networks_attr()” and “res_networks_attr” functions. Edge correlations were visualized Gephi (v0.9.2), with positive correlations shown in blue and negative correlations in orange. Networks were exported as “gexf” files. To assess network fragmentation, modularity based on edge weights was calculated in Gephi using the “Modularity” tool from the “Statistics” panel^42,43^. Nodes importance was evaluated using betweenness centrality via the “Network Diameter” tool in Gephi, which computes the shortest paths between nodes^44^. Final network visualizations were refined and exported as Scalable Vector Graphics (SVG) using Inkscape (v1.3.2).

All high-throughput sequencing datasets (raw reads) have been deposited in the European Nucleotide Archive (ENA) at the European Molecular Biology Laboratory-European Bioinformatics Institute (EMBL-EBI) under the project accession number PRJEB91711.

## 3. Results and Discussion

### 3.1 Spatial Pollution profiles in the Aconcagua and Maipo Rivers

To address our ecological questions regarding cross-domain co-occurrence, we integrated physicochemical profiling (Figure 1), including pH, conductivity, temperature, dissolved oxygen, dissolved organic carbon, nutrients as nitrite, nitrate, and phosphate (Supplementary Table 3) with measurements of OMPs previously reported in the Aconcagua and Maipo river basins (Inostroza et al., 2025; Soriano et al., 2024). The river basins are characterised by a distinct pollution gradient across sites subject to varying water demands, effluent discharges from WWTPs, and urban and agricultural runoff (Supplementary Figure 1). In addition to the OMPs, we determined the concentrations of various metals, including aluminium, cadmium, copper, and lead, among others (Figure 1).

**Figure 1.**
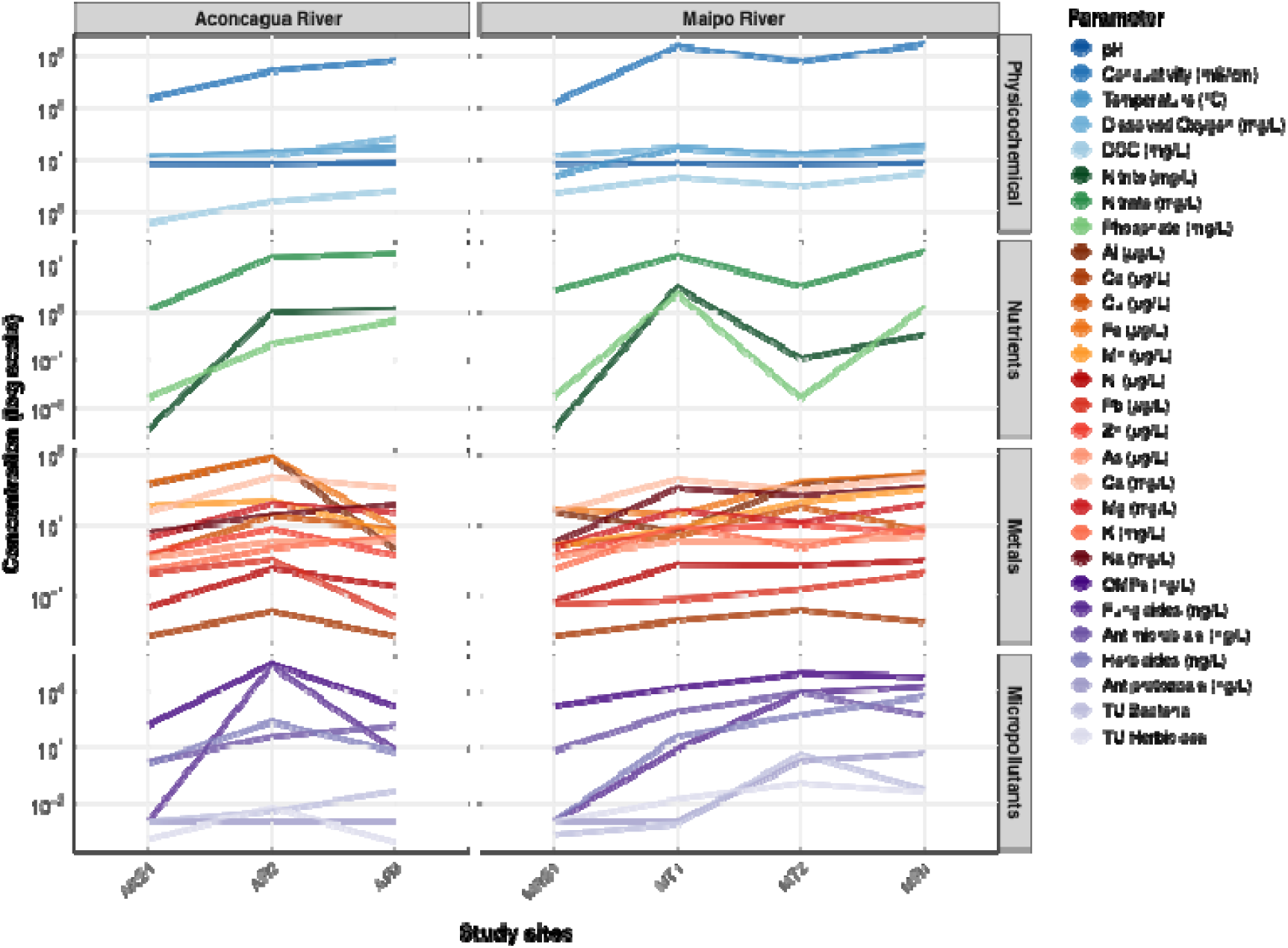
Environmental variables per river basin and across sites. TU Bacteria and TU Herbicides represent the OMP normalised concentrations of antimicrobial and photosynthesis inhibitor chemicals. (for details see section 2.2. in Methods).

We classified sampling locations into three pollution categories based on the measured physicochemical parameters, metals, and OMP concentrations (Soriano et al., 2024): reference (low pollution), rural (moderate pollution), and urban (high pollution). In the Aconcagua River, most measured variables demonstrated a longitudinal pollution gradient extending from the reference conditions at ARS1 to the most downstream sampling locations (AR3). Peak concentrations across several contaminant classes, including nutrients (nitrate, nitrite), metals, and OMPs (fungicides and antimicrobials), were evident at AR2, which is situated immediately downstream of major urban and agricultural areas (Figure 1). This spatial trend in contaminant profiles is consistent with previous studies characterising the pollution gradient within this region (Inostroza et al., 2024, 2023).

In the Maipo River, site MT2, located downstream of the main WWTP serving Santiago city, exhibited the highest concentrations of metals and the most diverse mixture of OMPs, including fungicides, antibiotics, and herbicides. The upstream reference site, MRS1, showed low levels of most pollutants, although it displayed elevated metal concentrations associated with the pre-Andean geology. The urban site MT1 (upstream of the WWTP) and the rural site MR1 were primarily characterized by nutrient enrichment and increased pH. These site-specific pollution profiles provided a basis to test the SGH in cross-domain microbial networks.

### 3.2 Network Fragmentation as a Hallmark of Community Destabilization

The predicted correlation patterns among bacteria, protists, and fungi, were consistent with the Stress Gradient Hypothesis (SGH): sites with elevated OMP loads exhibited a clear predominance of positive correlations (Table 1). In the Aconcagua River, the most polluted site (AR2) showed 100% positive correlations, while in the analogous site in the Maipo River (MT2) displayed 98.5% positive correlations. In contrast, rural sites with moderate nutrient enrichment but lower OMP levels showed a mixture of positive and negative correlations: 94.8% positive correlations at AR3 and 57.3% at MR1. Similarly, the urban site MT1, located upstream of a WWTP, had 58.5% positive correlations, comparable its rural counterpart MR1. Interestingly, even reference sites displayed most positive correlations–55.2% at ARS1 and 60.3% at MRS1–although negative correlations were more pronounced here than in the highly polluted sites.

**Table 1.** Networks topology information per study site at the Aconcagua and Maipo rivers.

|  | Aconcagua |  |  | Maipo |  |  |  |
| --- | --- | --- | --- | --- | --- | --- | --- |
|  | ARS1 | AR2 | AR3 | MRS1 | MT1 | MT2 | MR1 |
| Nodes | 149 | 107 | 179 | 136 | 122 | 152 | 161 |
| Edges | 8,108 | 1,725 | 5,729 | 5,939 | 4,644 | 4,931 | 9,940 |
| Positive correlations (%) | 55.29 | 100 | 94.83 | 60.3 | 58.55 | 98.5 | 57.39 |
| Negative correlations (%) | 44.71 | 0 | 5.17 | 39.7 | 41.45 | 1.5 | 42.61 |
| Modularity | 0.083 | 0.621 | 0.288 | 0.159 | 0.917 | 1.157 | 0.085 |
| Communities | 3 | 3 | 3 | 3 | 3 | 3 | 3 |

These findings suggest that facilitative interactions may be intrinsic characteristic of sediment microbiomes but become significantly more pronounced under chemical stress. Notably, differences between river basins were observed: while the Aconcagua reference site (ARS1) exhibited 55.2% positive correlations, increasing linearly along the pollution gradient, the Maipo reference site (MRS1) already had higher baseline of positive correlations (60.3%). A plausible explanation for this difference is that high metal concentrations, characteristic of the pre-Andean geology and consistent with large-scale copper mining activities in the areas (Cortés-Miranda et al., 2024), promote baseline facilitation within the microbial communities.

Overall, the prevalence of positive correlations was river-basin specific, yet consistently high in sites with elevated pollution. These results align with the SGH, which predicts an increase in facilitative interactions under heightener environmental stress (Hammarlund and Harcombe, 2019). In microbial communities, dominance of positive associations under such conditions is often interpreted as ecological cooperation–including resource sharing, syntrophy, or shared stress-tolerance mechanisms–facilitating persistence in adverse environments (Brenes-Guillén et al., 2022). Consistently, a Jaccard-based hierarchical clustering of overall community composition revealed that bacteria, protists and fungal assemblages tended to cluster by study site and taxonomic domain (Supplementary Figure 2). This result supported the robustness of the metabarcoding approach in capturing ecologically meaningful patterns that can inform subsequent network-based analyses.

### 3.3 Network Modularity and Fragmentation

Regarding network modularity, we hypothesised that sites featuring low pollution pressures (nutrients, metals, OMPs) would exhibit compact cross-kingdom networks. This was supported by low modularity values of 0.083 and 0.159 in the reference sites ARS1 (Aconcagua) and MRS1 (Maipo), respectively (Table 1). Conversely, cross-kingdom networks from polluted sites displayed significantly higher modularity (AR2=0.621 and MT2=1.157 in Aconcagua and Maipo, respectively), which is indicative of fragmentation (Hernandez et al., 2021). Interestingly, this fragmentation was observed in the most polluted site (AR2) and persisted in the downstream site AR3 of the Aconcagua River (Figure 2). In contrast, while networks in the Maipo River were also fragmented at polluted sites (MT2, downstream of the WWTP), they returned to a compact configuration in downstream areas (Figure 3). This difference may reflect higher baseline levels of positive correlations and naturally elevated metal concentrations in the Maipo basin, which could potentially buffer the destabilizing effects of anthropogenic pollution and maintain more integrated, compact network structures in downstream sites.

**Figure 2.**
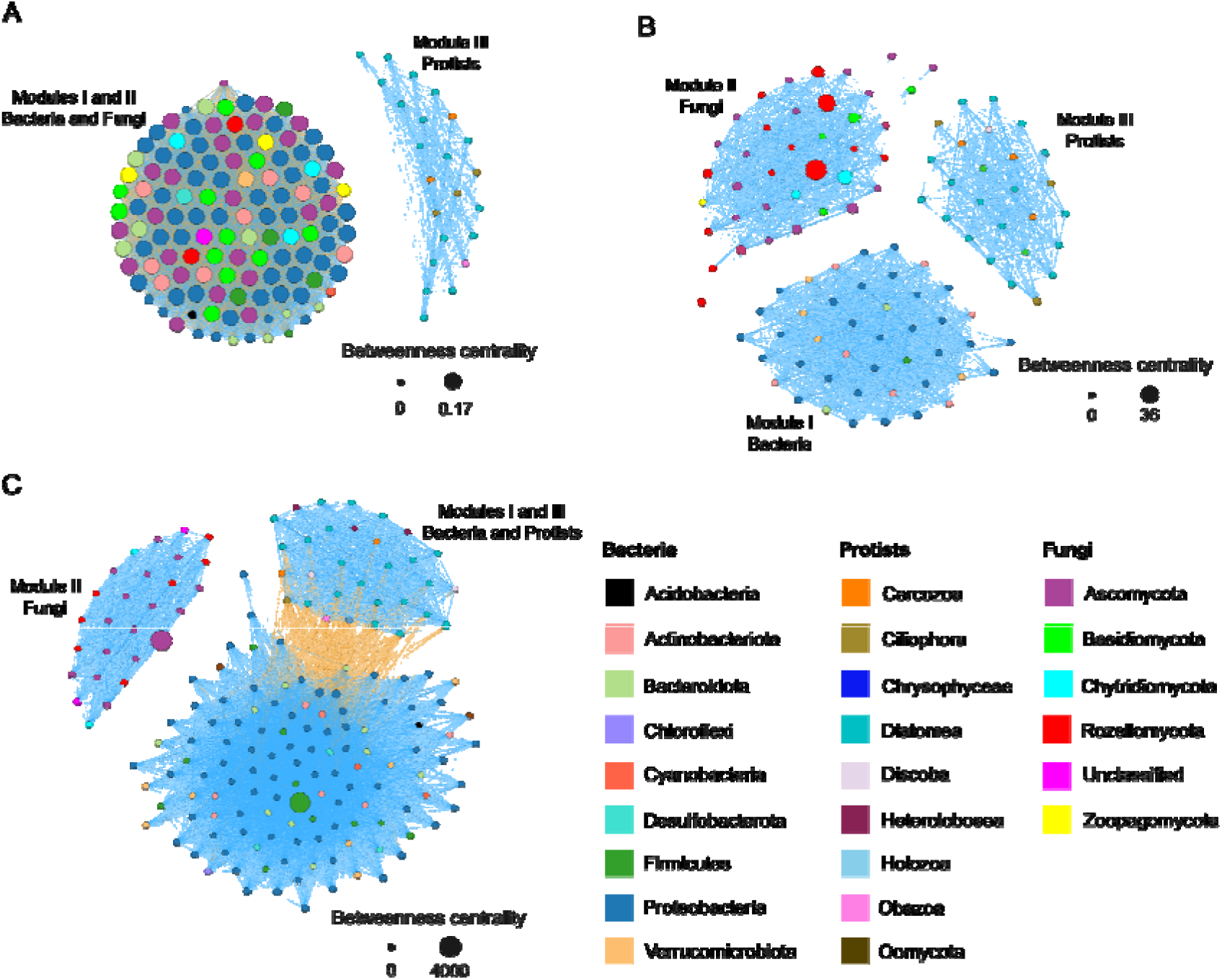
Co-occurrence networks and modules of genotypes classified as bacteria, protists and fungi detected at the study sites in Aconcagua River. A: Reference Site ARS1; B: Site AR2; C: Site AR3. Edges representing positive correlations are shown in blue and edges representing negative correlations are shown in orange.

**Figure 3.**
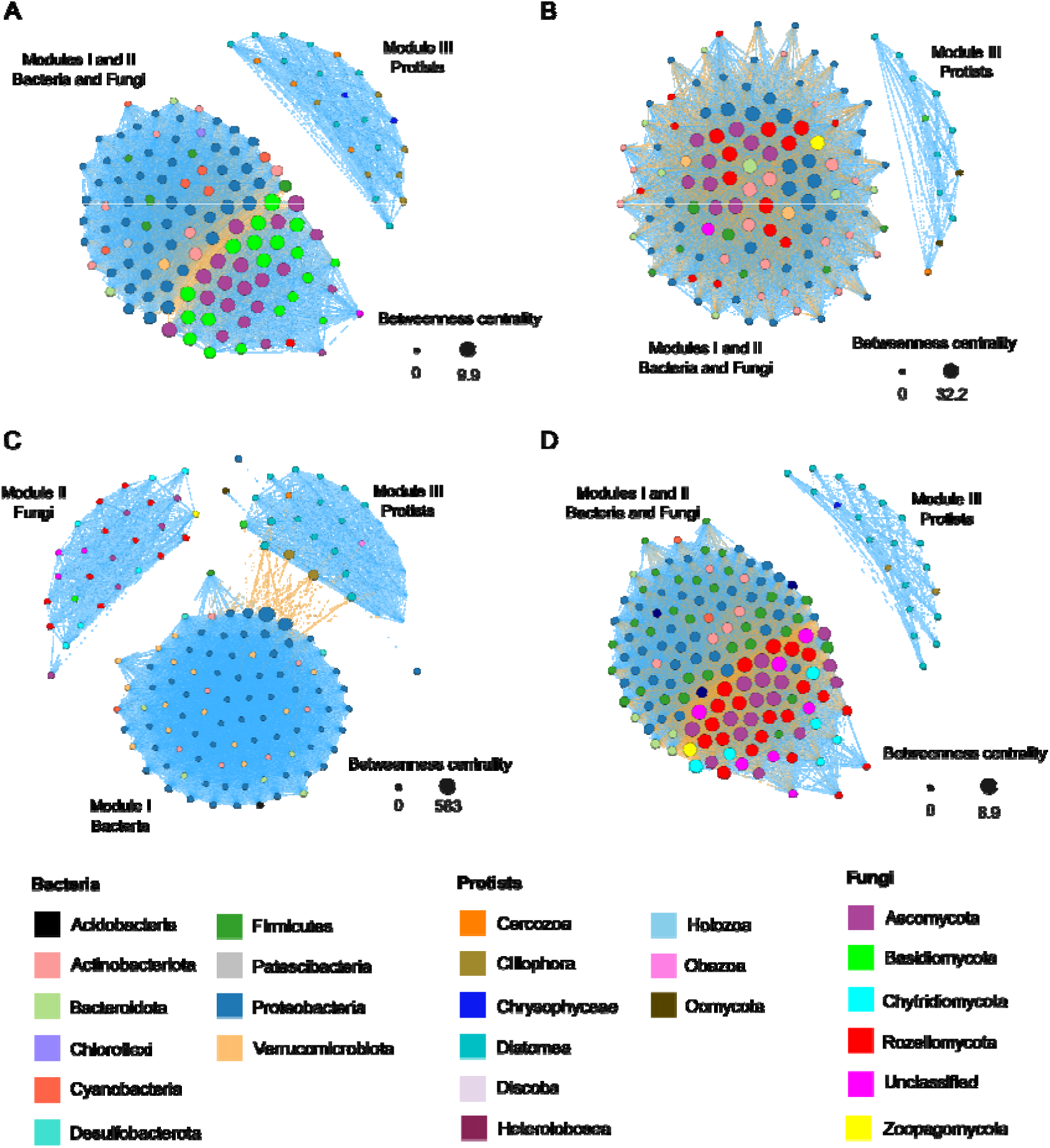
Co-occurrence networks and modules of genotypes classified as bacteria, protists and fungi detected at the study sites in Maipo River. A: Reference site MRS1; B: Site MT1; C: Site MT2; D: Site MR1. Edges representing positive correlations in blue and edges representing negative correlations in orange.

We observed that reference sites (Figure 2A and 3A) showed bacterial and fungal modules to be closely interconnected, forming cohesive node clusters. Conversely, polluted sites exhibited a pronounced separation between bacterial and fungal modules (Figure 2B and 2C, 3B and 3C). At the sites with the highest OMPs pollution (AR2 and MT2), network fragmentation intensified, marked by the emergence of isolated modules dominated by bacteria, protists or fungi. As shown in Table 1, these modules are still identified as separate communities, independent of the physical proximity between them. This fragmentation of the networks into isolated taxonomic clusters suggests potential niche partitioning or antagonistic interactions rather than cooperation (Weisse, 2017). At the rural site AR3 (Figure 2), bacterial and protist modules were clearly separated and negatively correlated–a pattern also observed at MT2 (Figure 3), a site located downstream of the country’s largest municipal WWTP. In these rural sites in both basins, fungi formed an isolated module, potentially reflecting the primary effect of pollution on microbial cross-kingdom interactions by disrupting the cohesion of bacteria-fungi modules observed at reference sites.

These clustering patterns are challenging to compare due to limited studies conducting cross-kingdoms networks in urban river sediments. Nonetheless, the observed shift toward taxonomic isolation and reduced inter-domain connectivity may indicate a stress-induced disruption of microbial interaction networks, consistent with observations from other polluted freshwater ecosystems (Hernandez et al., 2021). The disruption of microbial networks under pollution stress could have critical implications for ecosystem functioning due to the ecological role of microbial consortia (e.g., nutrient cycling, organic matter decomposition, and contaminant degradation) (Marie Booth et al., 2019; Miao et al., 2022). The loss of cross-domain interactions and the increase in modularity–such as observed at AR2–are likely to reduce functional redundancy and network robustness, thereby increasing vulnerability to additional stressors, including climate variability, hypoxia, and hydrological extremes (Faust and Raes, 2012; Hu et al., 2017).

### 3.4 Keystone Taxa and Domain-Specific Interaction

The taxonomic classification revealed that bacterial communities were dominated by Alpha- and Gammaproteobacteria, Bacteroidota and Actinobacteria in both river basins and across the distinct sampling sites (Supplementary Figure 3); These taxa are known key heterotrophs and chemoorganotrophs involved in carbon cycling, organic matter degradation, and nutrient transformations, particularly nitrogen and sulphur cycling (Agler et al., 2016; Piccardi et al., 2019). Protist communities were mainly composed by Ochrophyta, Ciliophora, and Cercozoa (Supplementary Figure 4); These groups encompass diverse autotrophic and heterotrophic lineages that regulate microbial food webs dynamics, often acting as bacterivores and influencing biogeochemical fluxes (Rizzatti et al., 2017). For instance, Ciliophora exhibit diverse feeding strategies, including predation and parasitism (Nguyen et al., 2023). Furthermore, fungal communities were characterised by Dothiomycetes, Sordariomycetes, and Tremellomycetes (Supplementary Figure 5). These groups are typically involved in the decomposition of organic matter, particularly plant-derived polymers such as lignin and cellulose (Dumack et al., 2022; Pohl et al., 2021), contributing to carbon turnover in sediments and potentially engaging in competitive or synergistic interactions that influence nutrient cycling (García-Carmona et al., 2021). While these patterns provide a baseline view of community composition, keystone taxa identified through betweenness centrality were not necessarily the most abundant community members.

We identified nodes with higher number of correlations using the betweenness centrality, a metric depicting a taxon’s topological importance within the network (Geesink et al., 2024). Among bacterial nodes, Proteobacteria were consistently central in reference sites (Figure 2A and 3A), suggesting their foundational role in maintaining network stability. However, with increasing pollution, these groups lost hub status, particularly in the Aconcagua River (Figure 2B), indicating sensitivity to environmental change. Conversely, at the urban site MT1 (upstream of the WWTP), Proteobacteria nodes became highly correlated, not only to other bacterial taxa, but also to protists nodes, primarily via negatively correlations (Figure 3B). At the most downstream polluted site, MT2 (downstream of the WWTP), Proteobacteria retained a central role, exhibiting high betweenness centrality and correlating with both bacterial nodes and to fungal nodes (e.g., to Rozellomycota). At the rural study site MR1, unlike its Aconcagua counterpart AR3, Proteobacteria showed close, predominantly negative correlations with fungal nodes (Figure 3D). Conversely, Firmicutes emerged as a central node in the Aconcagua rural site AR3, and as several highly connected nodes upstream of a WWTP (Figure 3B), reflecting their known opportunistic behaviour in stressful environments (Verweij et al., 2020). These patterns mirror shifts in topological roles observed in other microbiomes under stress, such as plant-associated communities, where disturbance in abiotic factors as temperature and type of habitat and climate often drives a turnover in few central bacterial taxa (Muteeb et al., 2023).

Protist communities showed compositional stability but diverged topologically across sites. In reference locations, protists often formed isolated modules without significant cross-domain interactions (Figure 2A and 3A). However, in rural sites and downstream of a WWTP, protists taxa–particularly Cercozoans and Ciliates–displayed increasingly negative correlations with bacterial nodes (Figure 2C). This suggests a shift toward intensified predator-prey or competitive interactions under chemically stressful conditions. Cercozoa, known for their trophic versatility and adaptability, have been proposed as bioindicators of anthropogenic pollution due to their frequent detection in WWTPs effluents and associations with both beneficial and opportunistic bacteria (H.R. Branco et al., 2023; Xie et al., 2022). Similarly, Ciliophora exhibited negative associations with bacterial taxa at a site with high nutrient loads (Figure 3D), supporting a top-down control mechanisms that may emerge as part of resilience and recovery in stressed ecosystems. Another important negative pattern included the negative correlation between bacteria and Diatomea (Figure 2C). Such antagonistic dynamics have been documented in benthic systems, where diatoms and bacteria share similar ecological niches, especially in nutrient deprived environments. Overall, these interaction patterns underscore the role of protists as central players in the microbial loop, restructuring under pollution stress, with critical implications for nutrient cycling, bacteria turnover, and food web stability in freshwater sediments.

Fungal communities showed high heterogeneity in their response to pollution gradient. Ascomycota remained the most ubiquitous and central taxon across all sites, maintaining key interactions in reference, rural and urban environments. These patterns align with reports suggesting that Ascomycota display broad environmental tolerance, while certain genera may serve as disturbance indicators under urban pressures. In contrast, Rozellomycota– a group with largely uncharacterized ecological roles–gained topological prominence under stress, forming highly connected nodes in AR2 and MT2. At the urban site AR2 in Aconcagua, Rozellomycota is central, however within an isolated fungal module. In contrast, this group was depicted as central in the urban site, upstream to the WWTP in Maipo River (MT1). Their emergence as central taxa in urban and nutrient-elevated rural sites points to a potential sentinel role in the sediment ecosystem, echoing recent studies that propose fungi as early indicators of ecological degradation. In contrast, the group Basidiomycota showed importance mainly at reference sites, suggesting a potential indication of more pristine conditions.

Collectively these patterns reveal how pollution gradients reshape the topology of cross-domain microbial networks and weaken cross-domain connectivity, with potential consequences for sediment ecosystem functioning. Such shifts potentially disrupt ecologic complementarity, especially across microbial domains, where protists and fungi may rely on bacteria syntrophy or substrate provisioning.

### 3.5 Limitation and future research

A key limitation of our approach is that co-occurrence networks are based on correlation analyses, which do not necessarily reflect direct ecological interactions or establish causality. Furthermore, DNA-based community profiling captures presence of organisms rather than their activity, which limits the resolution of ecological functional dynamics. To overcome these limitations, future efforts should integrate methods such as metatranscriptomics, functional metagenomics, metabolomics, or stable isotope probing to validate inferred interactions and clarify specific functional roles in comprehensive assessments. For rapid and cost-effective monitoring, cDNA metabarcoding could also be applied to provide an overview on the active– and potentially interacting– microbiome in the studied environments.

Moreover, while we included a wide array of chemical stressors, establishing a mechanistic understanding of their role, especially concerning OMPs, remains challenging under complex real-world conditions. Therefore, the observed network shifts must be tested under controlled experimental conditions. Controlled microcosms using single chemical and synthetic mixtures in a factorial design would help to dissect the complex natural dynamic observed in this study and isolate the impact of specific stressors (e.g., metals vs OMPs).

Finally, although the use of the correlation approach (SparCC) is validated for studies on compositional cross-domain datasets, emerging methods such as GLASSO-Split, may further improve network resolution and should be explored in future comprehensive cross-domain biodiversity studies.

## 4. Conclusion

Our findings revealed that anthropogenic pollution significantly restructures microbial interaction networks in freshwater sediments, characterised by distinct structural patterns along environmental gradients. We found a high degree of network fragmentation and isolation among bacterial, protist, and fungal communities in heavily impacted sites (high modularity), contrasting with the tighter bacteria-fungi coupling observed in less disturbed reference environments. This structural deterioration is interpreted as a loss of cross-domain connectivity and functional redundancy. Furthermore, the prevalence of positive correlations across both basins consistently supported the SGH, indicating a shift toward facilitative interactions under chemical stress. Our results also unveiled river-basin specific differences, with the Maipo River showing a higher baseline of positive correlations, likely due to naturally elevated metal concentrations promoting intrinsic facilitation. Notably, under moderate pressure, we observed a potential competitive or top-down dynamic between protists and bacteria (negative correlations), suggesting intensified trophic interactions that actively shape microbial energy and nutrient flow in transitional urban contexts.

Together, these findings highlight the value of network topology (modularity) and correlation directionality as sensitive and promising metrics for assessing ecological stress responses and resilience in complex microbial communities. Given the foundational role of protist-bacteria and fungi-bacteria interactions in ecosystem functioning, these patterns offer crucial insights into the cross-domain responses to human-driven environmental gradients. To deepen our understanding of these dynamics, time-series data and finer spatial resolution are critical for assessing network resilience and potential recovery following episodic contamination or restoration efforts. Our outcomes underscore the ecological relevance of interaction-based diagnostics in biomonitoring and highlight their utility as sensitive indicators of anthropogenic stress in freshwater sediments.

## 5. Acknowledgements

We thank Prof. Dr. Michael Schaub for for insightful discussions regarding the use of modularity maximization. We also thank Vicente Villalobos and Oliver Alarcón for their valuable support during fieldwork. We thank Dr. Alvaro Villalobos Claramunt for insightful discussions on the parameter setting for network construction and the calculation of genotype correlations. This work was supported by the FRAM Centre for Future Chemical Risk Assessment and Management at the University of Gothenburg, Sweden.

## 6. Author Contributions

CRediT authorship contribution statement: Eduardo Acosta: Writing – original draft, Data curation, Formal analysis, Investigation, Visualization. Thomas Backhaus: Writing - Review & Editing, Funding acquisition, Werner Brack: Writing - Review & Editing. Pedro A. Inostroza: Conceptualization, Methodology, Validation, Investigation, Writing - Original Draft, Writing - Review & Editing, Supervision, Project administration.

---

**Supplementary Figure 1.**
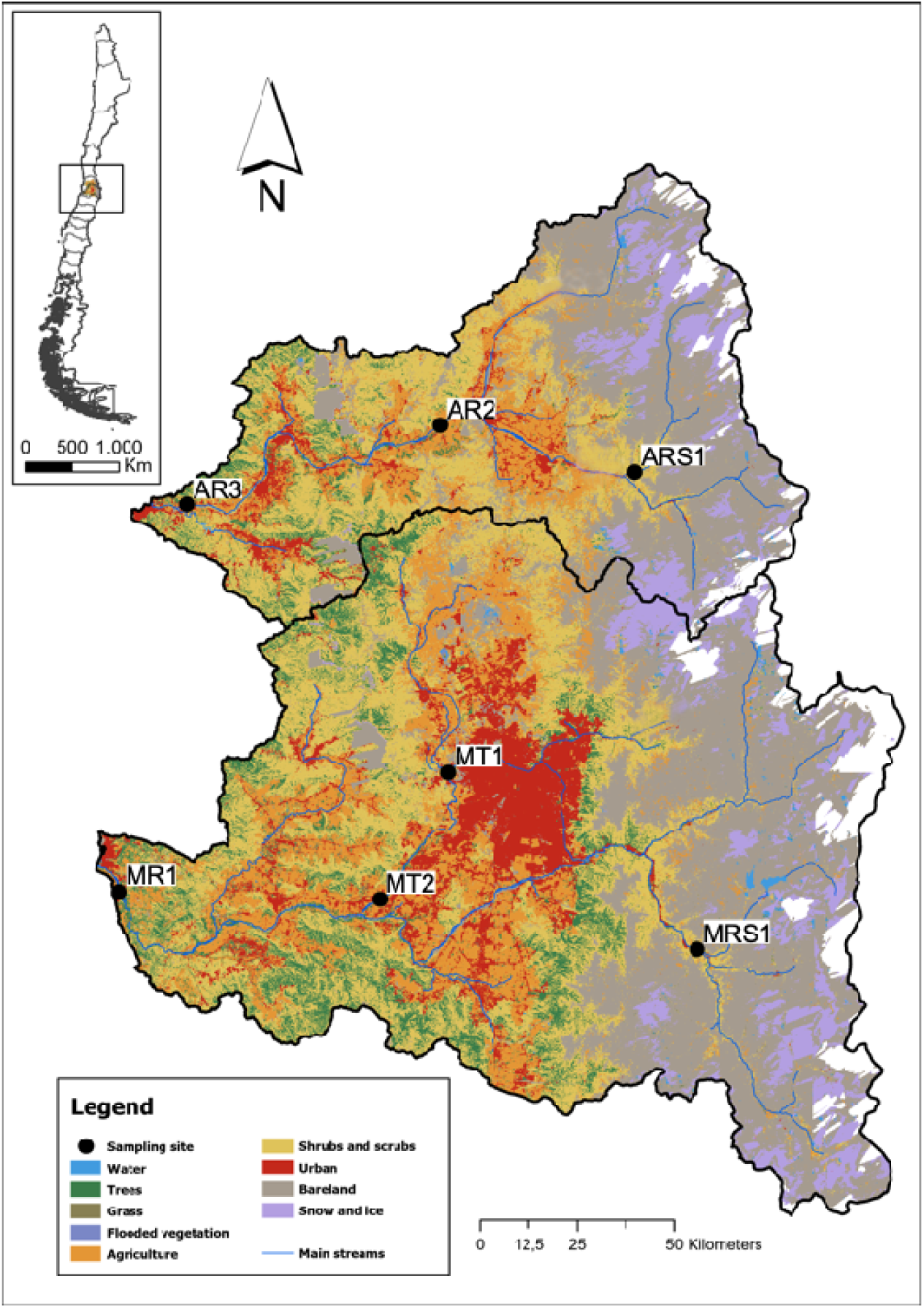
Map showing sampling sites at the Aconcagua (ARS1, AR2 and AR3) and Maipo (MRS1, MT1, MT2 and MR1) river basins. Figure is a modified version from Soriano et al., 2024.

**Supplementary Figure 2.**
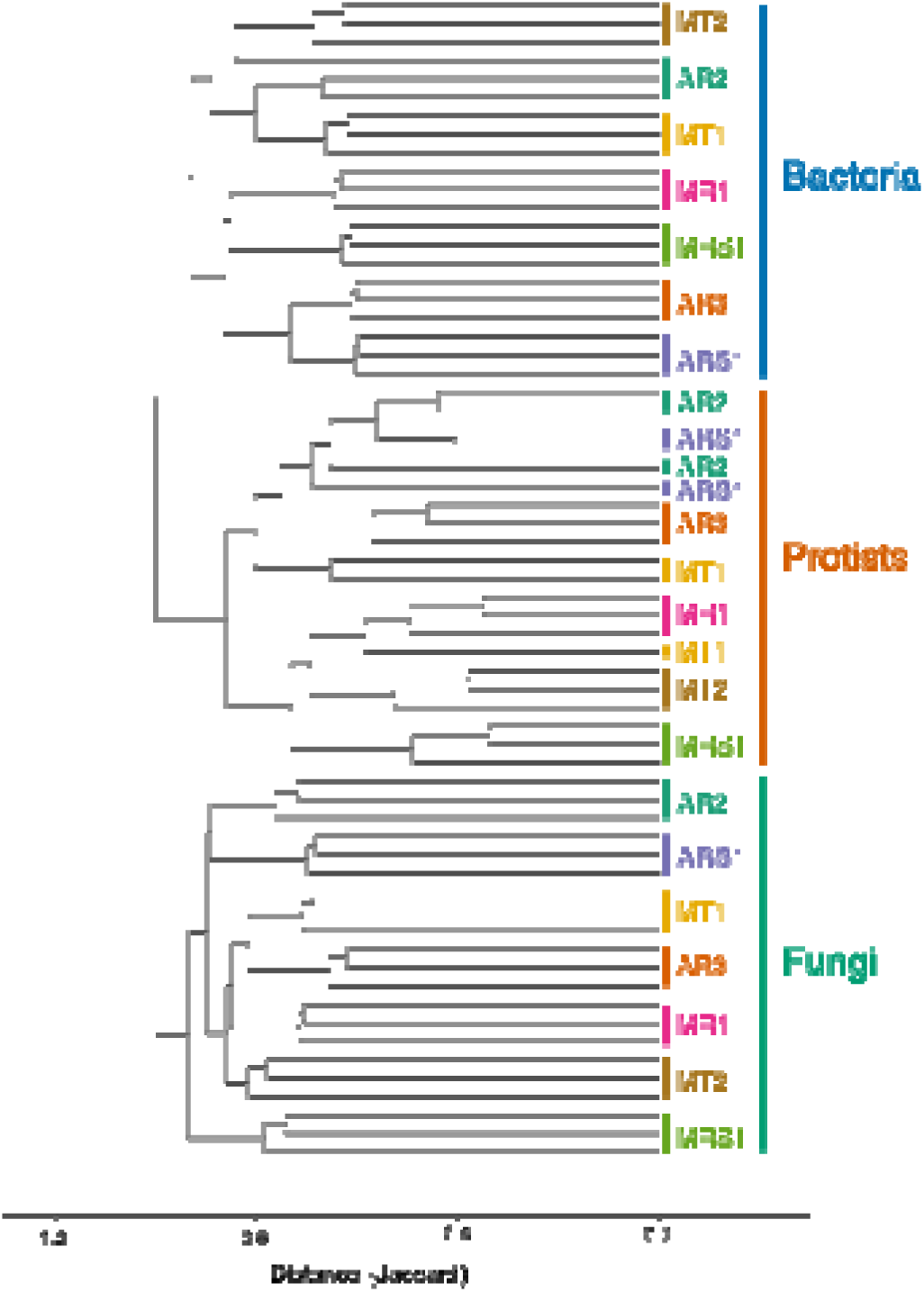
Cladogram based on the Jaccard distances among bacteria, protists and fungi detected in the study sites. (p < 0.001 for values >0.8).

**Supplementary Figure 3.**
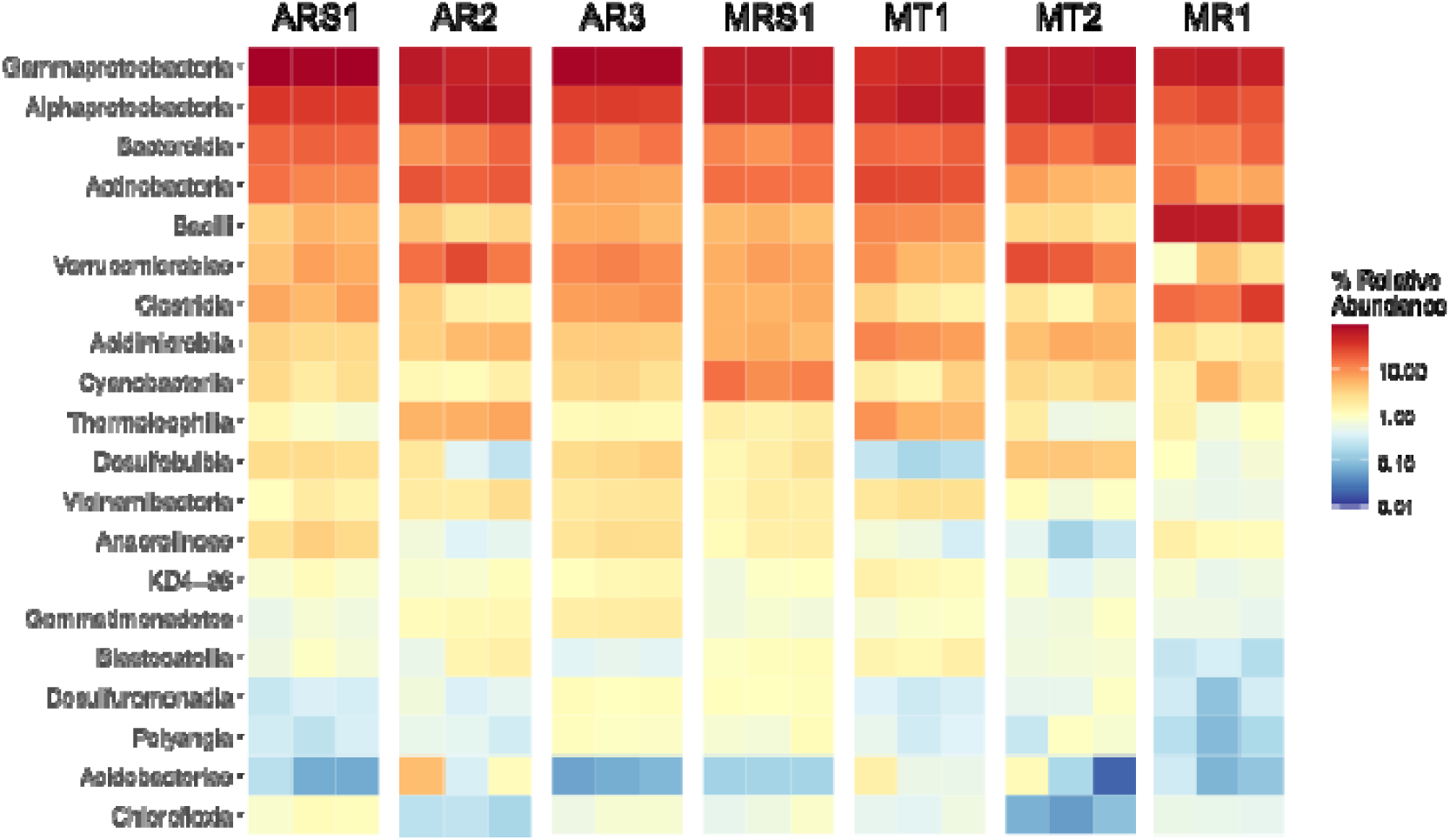
Top 20 bacterial groups classified at the Aconcagua (ARS1, AR2 and AR3) and Maipo (MRS1, MT1, MT2 and MR1) river basins.

**Supplementary Figure 4.**
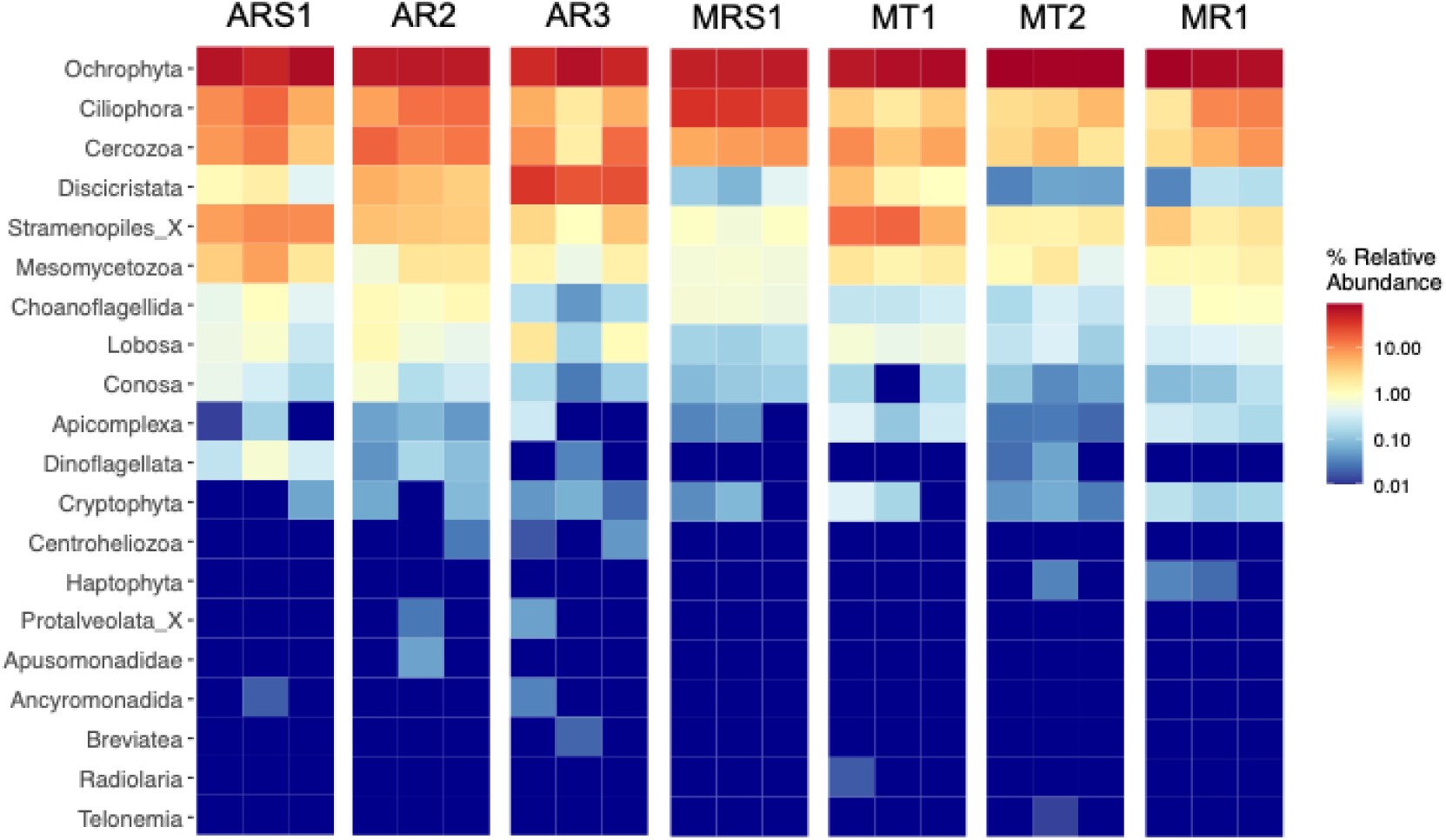
Top 20 protist groups classified at the Aconcagua (ARS1, AR2 and AR3) and Maipo (MRS1, MT1, MT2 and MR1) river basins.

**Supplementary Figure 5.**
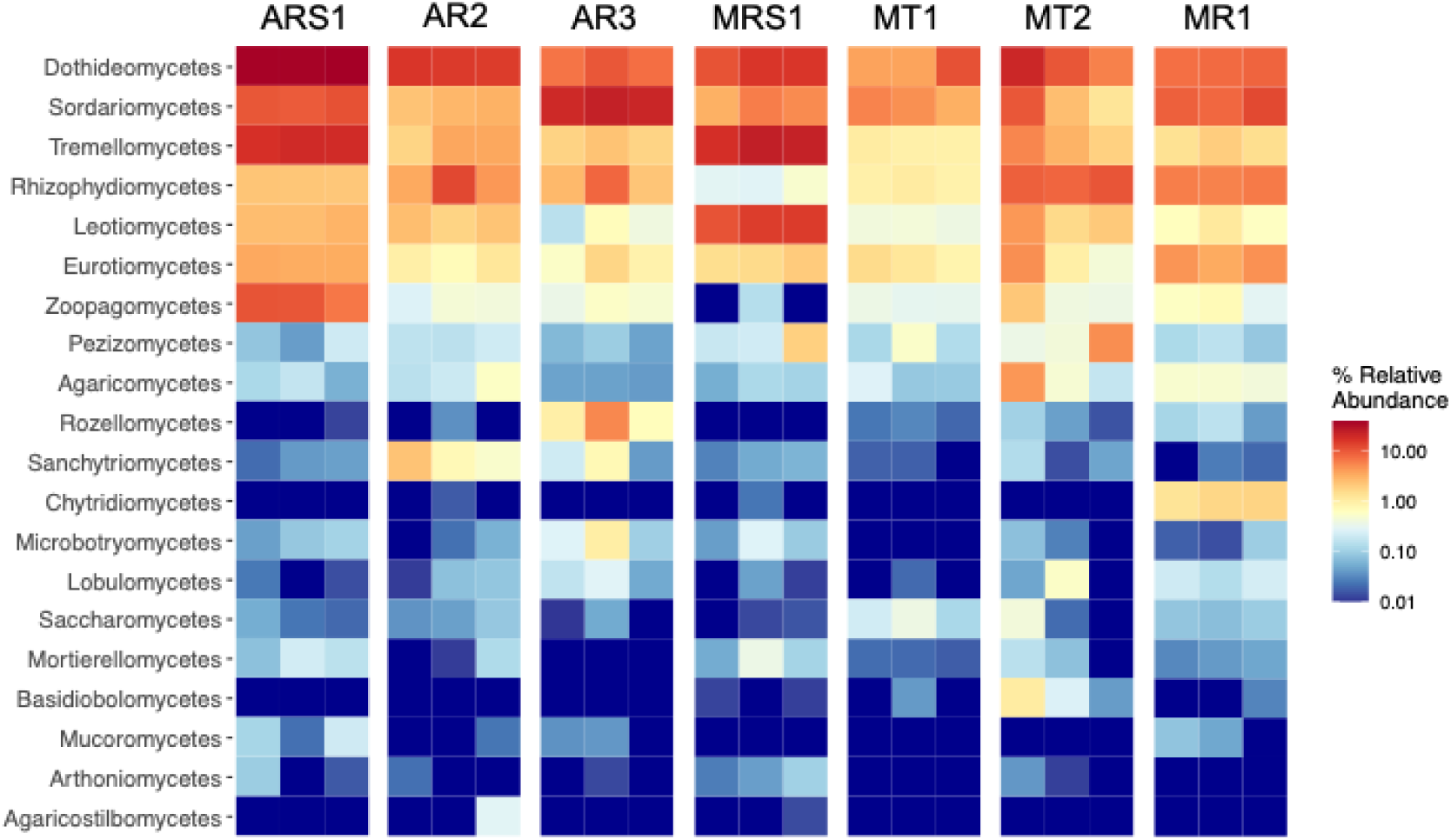
Top 20 fungal groups classified at the Aconcagua (ARS1, AR2 and AR3) and Maipo (MRS1, MT1, MT2 and MR1) river basins.

**Supplementary Table 1.**
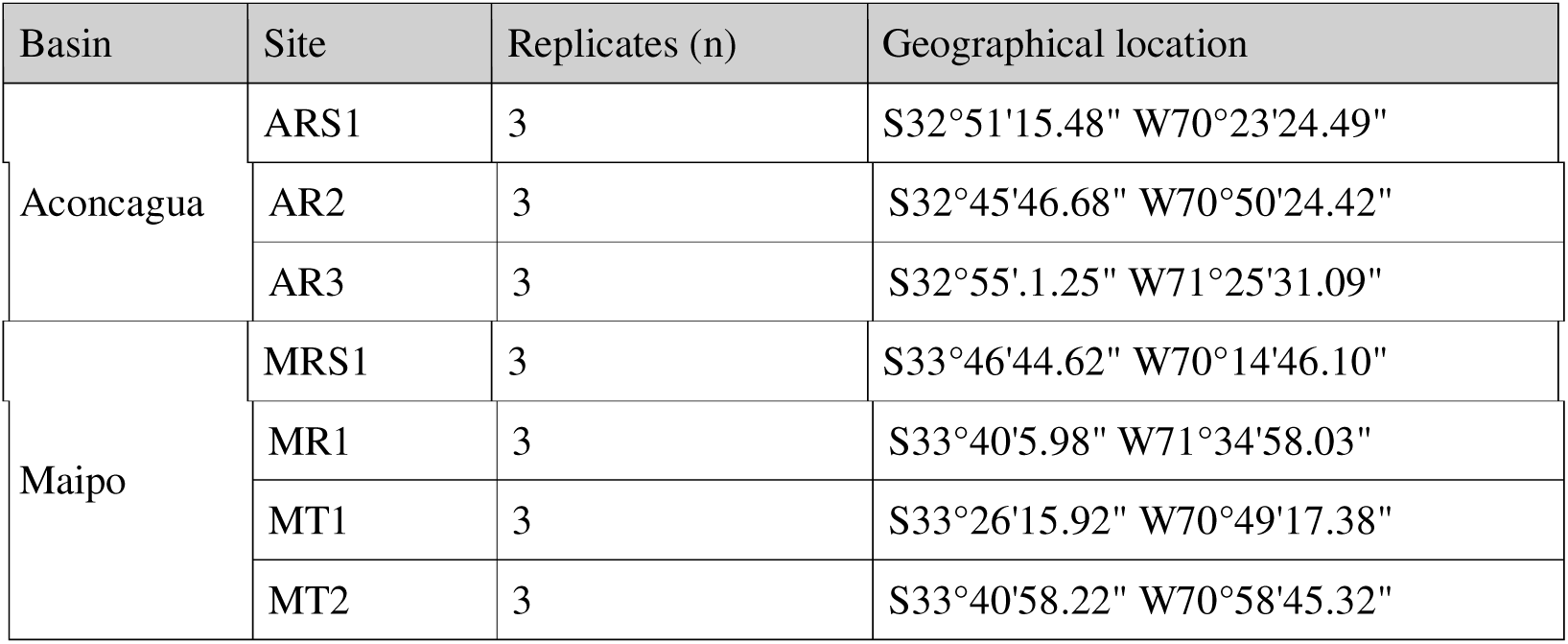
Sample information. Names, code, replicates, coordinates.

**Supplementary Table 2.** Concentrations and toxic units of OMPs measured at the study sites. Samples located upstream and downstream of a WWTP are referred as US-WWTP and DS-WWTP, respectively.

|  | Aconcagua |  |  | Maipo |  |  |  |
| --- | --- | --- | --- | --- | --- | --- | --- |
|  | ARS1 | AR2 | AR3 | MRS1 | MR1 | MT1 | MT2 |
|  | Reference | Urban/Agriculture | Rural | Reference | Rural | Urban (US-WWTP) | Urban (DS-WWTP) |
| Summed OMPs (ng/L) | 167.15 | 358,048.83 | 1,859.68 | 1,933.21 | 72,803.11 | 17,958.43 | 106,547.84 |
| Summed Fungicides (ng/L) | 0 | 283,352.3 | 18,810 | 0 | 20,365.12 | 45,816 | 9,668 |
| Summed Antimicrobial (ng/L) | 29,587 | 40.9 | 165.9 | 34,851 | 571.51 | 936.04 | 9,514.64 |
| Summed Herbicides (ng/L) | 14,977 | 266.6 | 35916 | 0 | 6,199 | 42.9 | 604 |
| Summed Antiprotozoal (ng/L) | 0 | 0 | 0 | 0 | 0 | 2 | 6 |
| Summed TU Bacteria | 0 | 0.0037 | 0.04606 | 0.0002 | 0.059 | 0.0007 | 4.74 |

**Supplementary Table 3.** Physic-chemical, nutrients, and metal concentrations.

|  | Aconcagua |  |  | Maipo |  |  |  |
| --- | --- | --- | --- | --- | --- | --- | --- |
| Parameters | ARS1 | AR2 | AR3 | MRS1 | MR1 | MT1 | MT2 |
| pH | 8.21 | 8.17 | 8.65 | 8.25 | 8.46 | 8.4 | 8.14 |
| Conductivity (mS/cm) | 148 | 506 | 796 | 125 | 1,654 | 1,501 | 763 |
| Temperature (°C) | 11 | 14 | 16 | 5 | 18.3 | 16.64 | 13.2 |
| Dissolved Oxygen (mg/L) | 12.06 | 12.09 | 25.78 | 12.19 | 15.6 | 17.04 | 11.45 |
| Nitrite (mg/L) | < 0.007 | 1.021 | 1.163 | < 0.007 | 0.347 | 3.433 | 0.11 |
| Nitrate (mg/L) | 1.134 | 14.777 | 18.191 | 2.871 | 20.168 | 16.049 | 3.568 |
| Phosphate (mg/L) | 0.016 | 0.216 | 0.696 | 0.017 | 1.31 | 2.514 | 0.016 |
| Dissolved organic carbon (mg/L) | 0.61 | 1.63 | 2.46 | 2.21 | 5.62 | 4.67 | 3.08 |
| Al (µg/L) | 145.03 | 829.72 | 1.94 | 23.27 | 268.41 | 6.66 | 152 |
| Cd (µg/L) | < 0.010 | 0.037 | < 0.015 | < 0.01 | 0.018 | 0.02 | 0.04 |
| Cr+6 (µg/L) | < 0.10 | < 0.10 | < 0.10 | < 0.10 | < 0.10 | < 0.10 | < 0.10 |
| Cu (µg/L) | 1.34 | 18.12 | 7.67 | 2.68 | 7.08 | 5.01 | 33.05 |
| Fe (µg/L) | 151.5 | 831.5 | 8.22 | 29.15 | 283.9 | 19.76 | 169.5 |
| Mn (µg/L) | 36.53 | 51.1 | 5.78 | 2.5 | 100.94 | 8.05 | 46.95 |
| Ni (µg/L) | 0.05 | 0.59 | 0.19 | 0.07 | 1.02 | 0.77 | 0.73 |
| Pb (µg/L) | 0.442 | 1.072 | 0.026 | 0.06 | 0.458 | 0.079 | 0.154 |
| Zn (µg/L) | 1.56 | 8 | 1.47 | 1.36 | 6.1 | 7.7 | 10.77 |
| As (µg/L) | 1.26 | 3.34 | 3.5 | 1.38 | 4.71 | 3.37 | 3.32 |
| Ca (mg/L) | 23.556 | 238.327 | 113.175 | 23.269 | 238.327 | 203.313 | 98.133 |
| Mg (mg/L) | 4.404 | 40.939 | 22.563 | 2.275 | 40.939 | 25.683 | 11.273 |
| K (mg/L) | 0.538 | 2.271 | 4.642 | 0.569 | 9.207 | 9.234 | 2.327 |
| Na (mg/L) | 6.259 | 18.852 | 39.683 | 3.244 | 119.671 | 109.005 | 65.397 |

